# Unusual Sulfur Requirements During Laboratory Growth of *Luteibacter*

**DOI:** 10.1101/149401

**Authors:** David A. Baltrus, A. Elizabeth Arnold

## Abstract

Many terrestrial bacteria are assumed to utilize sulfate transport and metabolism as a means for fulfilling cellular sulfur requirements. As such, many defined minimal media for bacterial growth under laboratory conditions contain sulfate as their sulfur source. Herein, an exception to this assumption is described as sulfate transport capabilities have been lost at least once in a lineage of *Luteibacter* associated with plants and fungi. However, a representative of this lineage (an endohyphal species, *Luteibacter* sp. 9143) can grow in minimal media when sulfur is supplemented with organic (cysteine and methionine) or inorganic (thiosulfate) compounds, and when co-cultured with its fungal host. A related strain of *Luteibacter* (UNC366Tsa5.1, isolated from the rhizosphere of Arabidopsis) potentially possesses more limited sulfur acquisition pathways than *Luteibacter* sp. 9143. These results highlight the surprising sulfur requirements of *Luteibacter*, which may be illustrative of close associations between these strains and eukaryotes, as well as a need for caution when inferring auxotrophies in a focal strain based on differential growth in minimal versus rich media.

**Importance:** Sulfate is often used as the sulfur source in minimal media. Here we show that some *Luteibacter* strains cannot utilize sulfate as a sulfur source, likely due to loss of genes encoding transport proteins. As sulfur requirements for *Luteibacter* can be met through co-culture with their fungal partner, this knowledge could provide a means to engineer better symbiotic relationships between bacteria and fungi that may be relevant for agriculture. Because growth in minimal media can be restored by supplementation with either cysteine or methionine, and in some cases only methionine, this result highlights how unexpected growth requirements could masquerade as auxotrophy for certain strains and conditions.

## Introduction

Sulfur enables critical physiological and chemical interactions within and between cells, such that it is an essential nutrient required across all life (1). Given the importance of sulfur for various biochemical processes, it is unsurprising that proteobacteria have evolved diverse mechanisms to obtain this sometimes limiting element. Even though these pathways range from scavenging sulfur from inorganic (e.g., sulfates and sulfides) or organic (e.g., cysteine, methionine, and sulfonates) sources, many proteobacteria are thought to rely on sulfate transporters CysAUW and CysP to internalize this element (1, 2). The prevalence of sulfate transport and metabolism has led to sulfur supplementation for a variety of defined growth media for aerobic bacteria (e.g., M9 and MOPS).

*Luteibacter* is a genus of Xanthomonadaceae (gammaproteobacteria) that often is found as a component of plant and fungal microbiomes (3, 4). Although little is known about their precise ecological functions, some *Luteibacter* strains act as plant growth promoting bacteria, with some producing phytohormones in partnership with endophytic fungi (5). *Luteibacter* sp. 9143 was isolated originally as an endohyphal bacterium from a foliar endophytic strain of *Pestalotiopsis* sp. 9143 (Ascomycota) (6). This *Luteibacter* grows readily in culture and has a facultative association with its fungal host (6).

When growth of *Luteibacter* sp. 9143 was characterized on different media, it was observed that this strain grew in rich media and when co-cultured with its fungal host in M9 media, but failed to grow in minimal media when grown axenically, suggestive of auxotrophy. A combination of selective supplementation, as well as screening of Biolog plates, demonstrates that this strain is not an auxotroph in the traditional sense. Instead, it cannot utilize sulfate as a sulfur source during laboratory growth. Further experiments show that *Luteibacter* sp. 9143 can grow in M9 media if supplemented with cysteine, methionine, or high levels of thiosulfate, or when co-cultured with its fungal host. Genomic evidence suggests that the requirement for sulfate-independent sulfur supplementation may be widespread in *Luteibacter,* with some strains possessing more limited sulfur scavenging capabilities than *Luteibacter* sp. 9143. These results suggest that strains of *Luteibacter* have evolved a unique metabolic niche for proteobacteria, potentially reliant on associations with hosts to obtain an essential element for growth.

## Methods

### Strains

The focal *Luteibacter* strain used across experiments in this study is DBL564, a rifampicin resistant isolate of *Luteibacter* sp. 9143 described in Arendt et. al 2016 (6). DBL966 is a rifampicin resistant isolate of *Luteibacter* sp. UNCMF366Tsu5.1 first described as associated with *Arabidopsis* by Lundberg et. al (3). This isolate was obtained from the Dangl Lab.

### Growth Curve Experiments

Populations of each *Luteibacter* strain were streaked from frozen stocks to Lysogeny Broth (LB) agar plates (10g of NaCl per L) supplemented with rifampicin at 50 ng/**μ**L and grown at 27^o^C. Prior to the start of each growth curve experiment (below), a single colony was picked and transferred to 2mL liquid LB media containing rifampicin and grown overnight. All liquid cultures were grown at 27^o^C on a rotating shaker (200rpm).

For fungal co-culture experiments, an isolate of *Pestalotiopsis* sp. 9143 was grown for 1 week in 3mL M9 media on a rotating shaker at 27^o^C and 200rpm. Mycelium was extracted from this culture and macerated by shaking with glass beads in 500**μ**L M9, after which 500**μ**L of additional M9 was added to this tube. A 100**μ**L volume of this resuspension was then added to 10mL M9 with *Luteibacter* sp. 9143 and 2mL aliquots from these master cultures were placed into 4 test tubes for each treatment. Fungal co-culture experiments were carried out twice independently, with negative controls treated as the experimental cultures except without the addition of fungi.

For bacterial growth curve experiments, 1mL of each culture was pelleted by centrifuge after overnight growth in LB media, washed twice in 10mM MgCl2, and resuspended in 1mL 10mM MgCl2. A subset of this resuspension was used to inoculate a master culture for each experiment as described below. All growth curve experiments were performed in standard M9 media with 4% glucose (i.e., M9). All M9 media contained 2mM MgSO4. The base M9 media was supplemented with additional chemicals where necessary (below). Supplements were added to a final concentration of either 100**μ**L or 10mM in M9 where indicated, except casamino acids (Difco), which were added at 0.1% w/v.

To begin a growth curve, 10mL of M9 media with supplementation was inoculated with 100**μ**L of *Luteibacter* and aliquoted in 2mL increments to test tubes. Bacterial count at the start of the experiment was measured by plating a dilution series of this inoculum on LB agar plates supplemented with rifampicin. Cultures were grown for at least 10 days, and cell counts measured by sampling a small volume (2**μ**L) of each culture and plating a dilution series on LB agar plates supplemented with rifampicin.

### Biolog Assays

For Biolog assays, an isolate of strain DBL564 was assayed by technicians at Biolog. Sulfur utilization was scored as positive if growth profiles in wells of assay exceeded average height threshold for both independent trials using proprietary software (Biolog, Inc).

### Data Availability

Growth data is available through Figshare at DOI: 10.6084/m9.figshare.5164864

### Phylogenetics and Genome-wide Comparisons of Sulfur Pathways

The complete genome sequence for *Luteibacter* sp. 9143 was described in a previous publication (7), and all further genomic comparisons took place through the Integrated Microbial Genomes (IMG) platform (http://img.jgi.doe.gov) developed by the JGI (8). Briefly, we inferred phylogenies using whole genome sequences for both strains investigated as well as closely related strains with available genomes. The phylogeny was first described in Baltrus et al. (2017) and shows a consensus maximum likelihood tree built from whole genome SNPs across the specified genomes using RealPhy (9). Strains marked with an asterisk were used as reference genomes. Genome annotations for all of these strains were then queried for annotations involving all KEGG pathways relevant for sulfur metabolism.

## Results and Discussion

### Co-culture with host fungus enables growth of *Luteibacter* sp. 9143 in M9 media without supplementation

In experiments designed to establish a co-culture system for *Luteibacter* sp. 9143 and its host (*Pestalotiopsis* sp. 9143), we observed that the supernatant of fungal-bacterial cultures in M9 media grew turbid over time under co-culture conditions, but not when bacteria were incubated axenically (data not shown). Subsequent growth curve experiments (Figure 1A) demonstrated that viable cell counts for *Luteibacter* sp. 9143 in supernatants increased over time during fungal co-culture but not in M9 alone. These observations strongly suggested that co-culture with its host fungus could provide, or provide access to, or more missing nutrients for *Luteibacter* sp. 9143 growth in M9 media.

**Figure 1.**
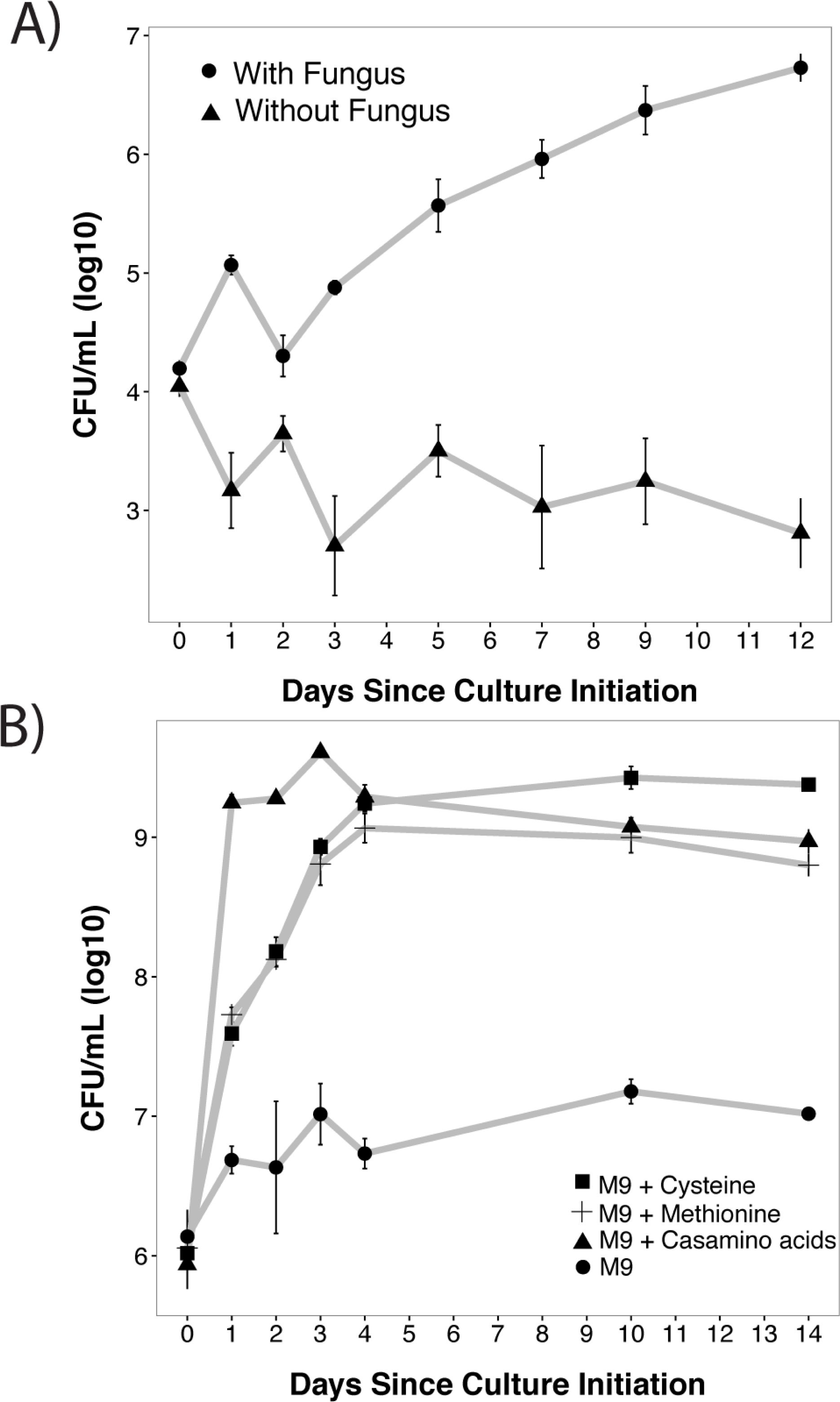
*Luteibacter* sp. 9143 Requires Sulfur Sources Other than Sulfate During Laboratory Growth. A) Representative growth curves of axenic *Luteibacter* sp. 9143 in M9 (triangles) or co-cultured with its host fungus, the foliar endophyte *Pestalotiopsis* sp. 9143 (circles). Four cultures were grown in each case, and bacterial population sizes in the supernatant were sampled and averaged at times indicated. B) Growth curves of axenic *Luteibacter* sp. 9143 in M9 media (circles), or supplemented with 100uM methionine (pluses), 100**μ**M cysteine (squares), or casamino acids (triangles). Each point shows the average from 8 replicates (over two different experiments) for each treatment combination. In all cases, error bars represent +/- one standard error.

### *Luteibacter* species grow in M9 media supplemented with additional sulfur sources

Further growth curve experiments confirmed that *Luteibacter* sp. 9143 was unable to grow axenically in base M9 media. However, this strain could grow if M9 was supplemented with casamino acids, suggesting that this strain was auxotrophic for at least one amino acid (Figure 1B).

Supplementation of M9 cultures with combinations of each amino acid within casamino acids suggested that either methionine or cysteine could support growth of *Luteibacter* sp. 9143 in M9 (data not shown). Growth curves (Figure 1B) demonstrate that *Luteibacter* can grow in M9 media if it is supplemented with either 100**μ**M cysteine or methionine. Growth was apparent after 10 days in each experiment regardless of treatment, such that day 10 chosen as a setpoint for further supplementation assays.

### Differential abilities of Luteibacter sp. *9143* to utilize organic and inorganic sulfur sources

Given that growth of *Luteibacter* sp. 9143 in M9 was sensitive to amino acids that contain sulfur, the production of which are often interconnected (1), the strain appears not to be a true auxotroph in the traditional sense. Unlike what is commonly assumed for culturable proteobacteria (1), *Luteibacter* sp. 9143 cannot utilize sulfate as a sulfur source.

Following up on these experiments, a variety of other sulfur sources were tested for their ability to supplement *Luteibacter* growth in M9 media. Serine was included because its structure is similar to cysteine. None of the assayed compounds, other than methionine and cysteine, could support growth of *Luteibacter* at 100**μ**M (Figure 2A). However, sodium thiosulfate supported growth when supplemented at 10mM (Figure 2B). Thus *Luteibacter* sp. 9143 can utilize organic sulfur sources like cysteine or methionine for growth under natural conditions, but can use thiosulfate if concentrations are high enough.

**Figure 2.**
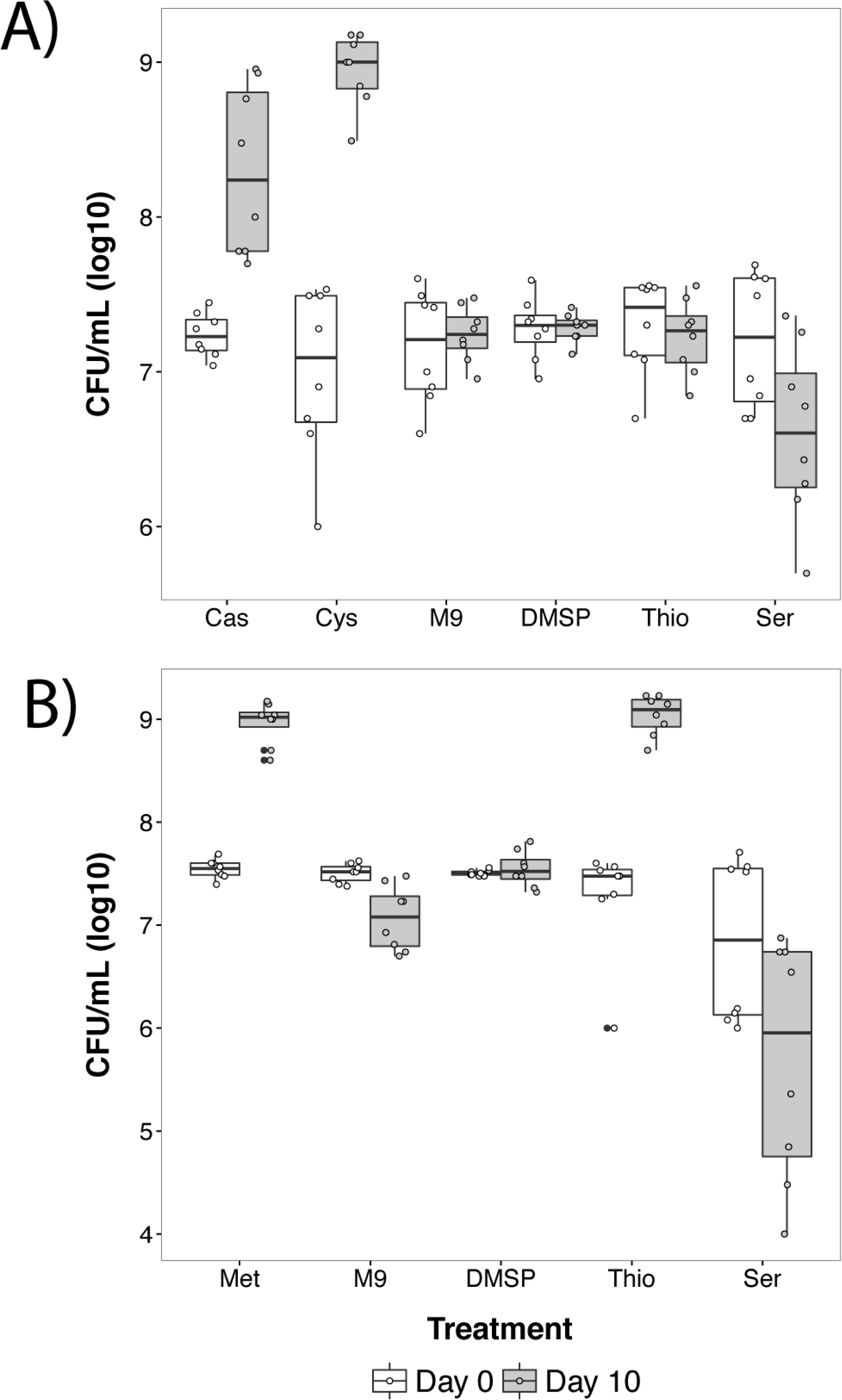
Cysteine, Methionine, and Thiosulfate Can Be Utilized as Sulfur Sources by *Luteibacter* sp. 9143. Plots show each data point for *Luteibacter* sp. 9143 cultures sampled at day 0 (white plot) and the same cultures sampled at day 10 (grey plot) in M9 supplemented with a variety of sulfur sources or serine. In each case, population sizes were sampled for each culture on both days. A) Cultures supplemented at 100μM with 0.1% w/v casamino acids (Cas), cysteine (Cys), DMSP, thiosulfate (Thio), serine (Ser), or unsupplemented (M9). B) Cultures supplemented at 10mM, with the only difference in treatment being methionine (Met) instead of cysteine. Error bars represent +/- one standard error.

The ability of *Luteibacter* sp. 9143 to utilize a variety of sulfur sources was assayed independently using Biolog phenotype array plates (Table 1 and Supplemental file 1). Data from two independent Biolog assays supported growth curve data, in that they showed that *Luteibacter* sp. 9143 could utilize both cysteine and thiosulfate as sulfur sources. The assays also showed that this strain could utilize Djenkolic acid, lipoamide, and lanthionine. In contrast to growth curve data, Biolog assays did not reliably support the ability of *Luteibacter* sp. 9143 to utilize methionine as a sulfur source (see report, on Figshare, at DOI: 10.6084/m9.figshare.5134852). We note, however, that activity of *Luteibacter* in multiple alternative sulfur sources does appear to be higher than that in sulfate according to the Biolog data. However, these substrates didn’t exceed the “average height” threshold according to Biolog’s proprietary software and therefore weren’t scored as positive growth.

**Table 1.**
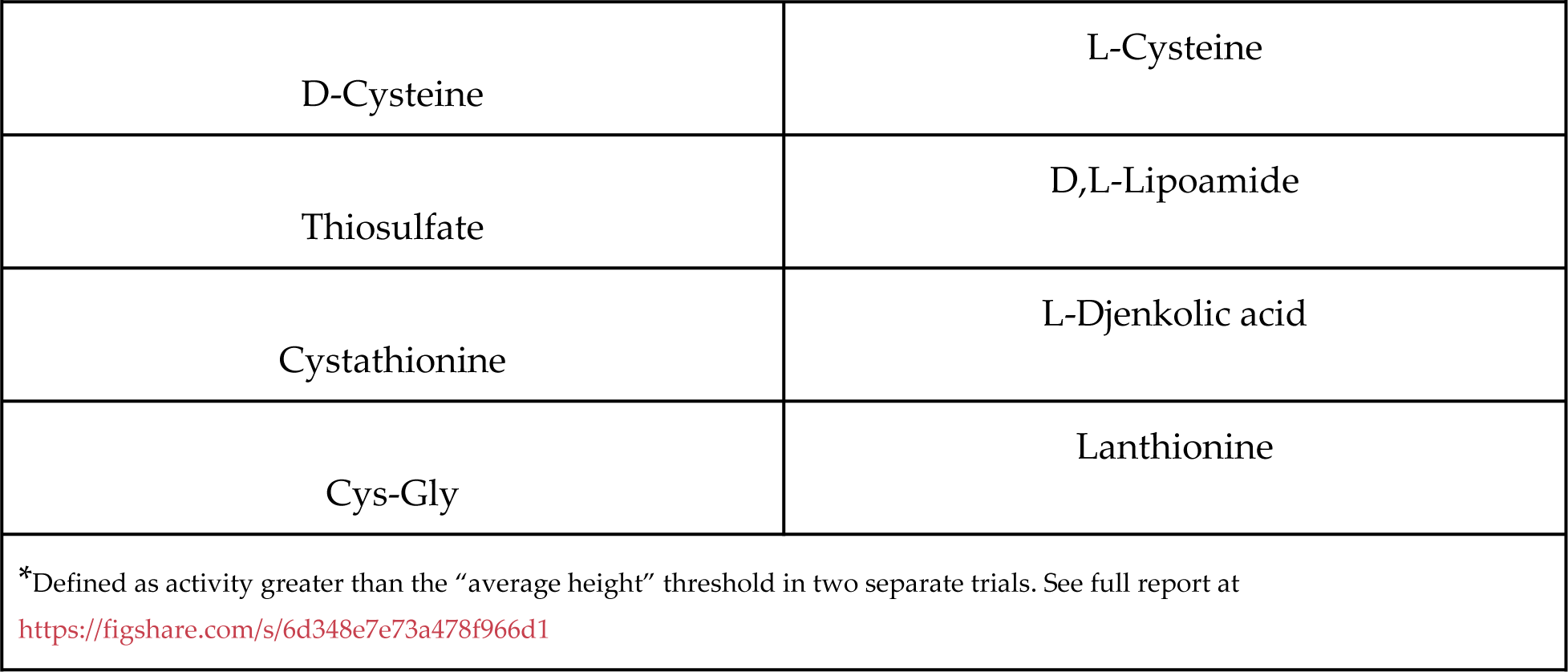
Sulfur Compounds that Repeatedly* Supported Growth of Luteibacter in Biolog Assays

### Genomic Comparisons Suggest that a Clade of *Luteibacter* Has Lost the Ability to Import Sulfate

JGI’s IMG server was used for pathway function analysis across *Luteibacter* genomes with genomes for *Dyella* and *Xanthomonas* as outgroups. The main genes responsible for sulfate transport in proteobacteria (*cysAUW* and *cysP/sbp*) are present in both outgroups and a subset of *Luteibacter* species, but are absent from the genomes of all strains found as endohyphal bacteria (Figure 3). These genes also are absent from the genomes of a clade of *Luteibacter* strains isolated from *Arabidopsis* roots. The most parsimonious interpretation of this pattern is that *cysAUWP* were lost independently from both clades of *Luteibacter*, or that they were lost in an ancestor of *Luteibacter* and reacquired by strains including *L. rhizoxinicus*. Either scenario would require two independent events (i.e., two losses of *cysAUWP* or one loss and one gain of *cysAUWP*).

**Figure 3.**
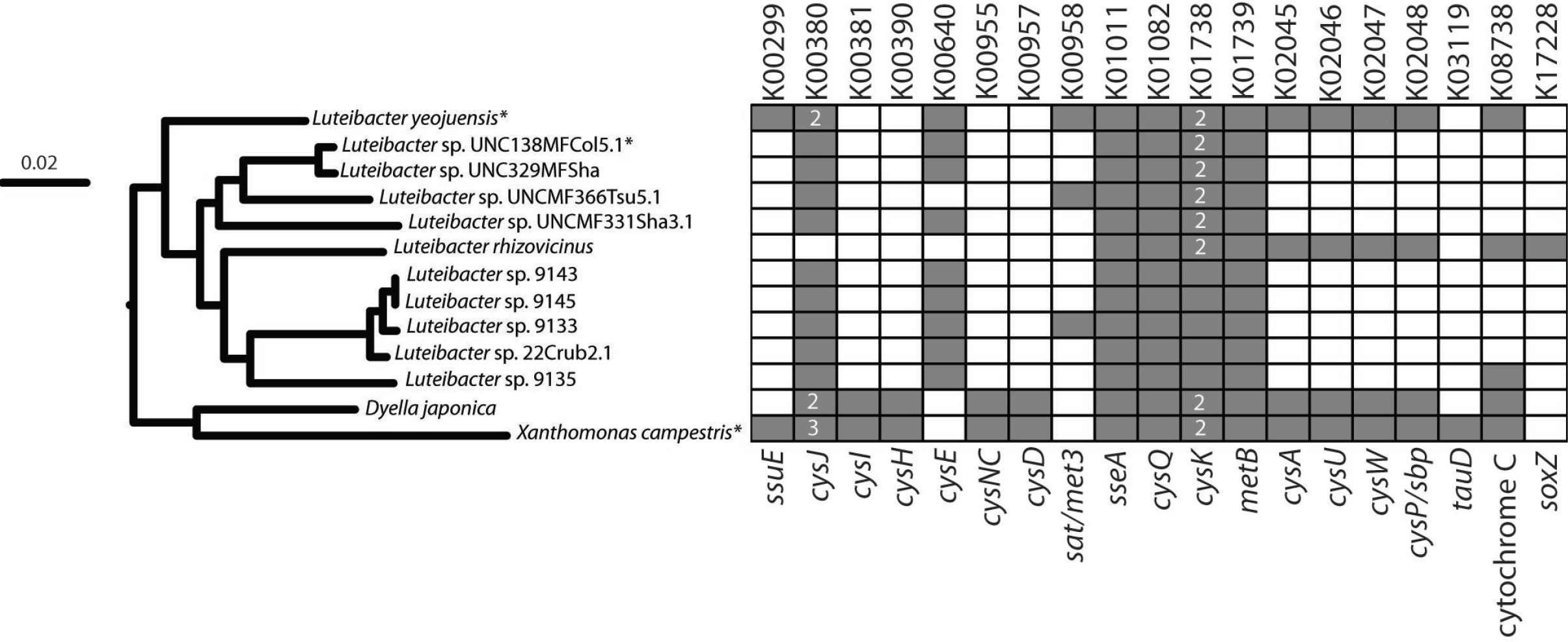
Multiple *Luteibacter* Strains Have Lost Ability to Transport Sulfate. A maximum likelihood phylogeny built using whole genome sequences from *Luteibacter* strains and related species, with methods explained in (7). Phylogeny was built in RealPHY using genomes listed here with an asterisk as references. Box on right shows presence (grey) or absence (white) of genes involved in sulfur utilization from the listed genomes as annotated in IMG. KEGG classifiers for each gene are shown at top of each column while gene names or descriptors are shown at bottom. Genes involved in sulfur acquisition, but which have no representatives in the queried genomes, are not shown in the matrix. Numbers inside boxes indicate that multiple copies of this gene(s) are found within the genome.

Genome comparisons suggested that other *Luteibacter* strains also would require a sulfur source other than sulfate for growth (Figure 3). Therefore, growth requirements of a rifampicin resistant mutant of *Luteibacter* sp. UNCMF366Tsu5.1 were assayed in M9 media alone and supplemented with sulfur sources shown to promote growth of *Luteibacter* sp. 9143.

Surprisingly, although *Luteibacter* sp. UNCMF366Tsu5.1 was able to grow in M9 supplemented with methionine, it was not able to grow in M9 supplemented with cysteine or thiosulfate (Figure 4). Two alternative, and not necessarily mutually exclusive, explanations could account for this differential growth behavior. First, transport and sulfur utilization for methionine could occur independently from that of cysteine and thiosulfate in these strains, and the latter could be missing from *Luteibacter* sp. UNCMF366Tsu5.1. Second, *Luteibacter* sp. UNCMF366Tsu5.1 may be more sensitive to reactive effects of cysteine and high levels of thiosulfate. Consistent population sizes (rather than cell death) of *Luteibacter* sp. UNCMF366Tsu5.1 in M9 with cysteine supports the first possibility and argues against the second, but a definitive answer cannot be reached from these data alone. Differential growth of *Luteibacter* strains also argues argues against contamination as the reason *Luteibacter* sp. 9143 could grow in 10mM but not 100**μ**M thiosulfate. It should be noted that, if *Luteibacter* sp. UNCMF366Tsu5.1 was assayed before *Luteibacter* sp. 9143, a natural conclusion would be that this strain was a methionine auxotroph. While this may be true, a more likely possibility is that *Luteibacter* sp. UNCMF366Tsu5.1 has a more limited sulfur scavenging capability than *Luteibacter* sp. 9143.

**Figure 4.**
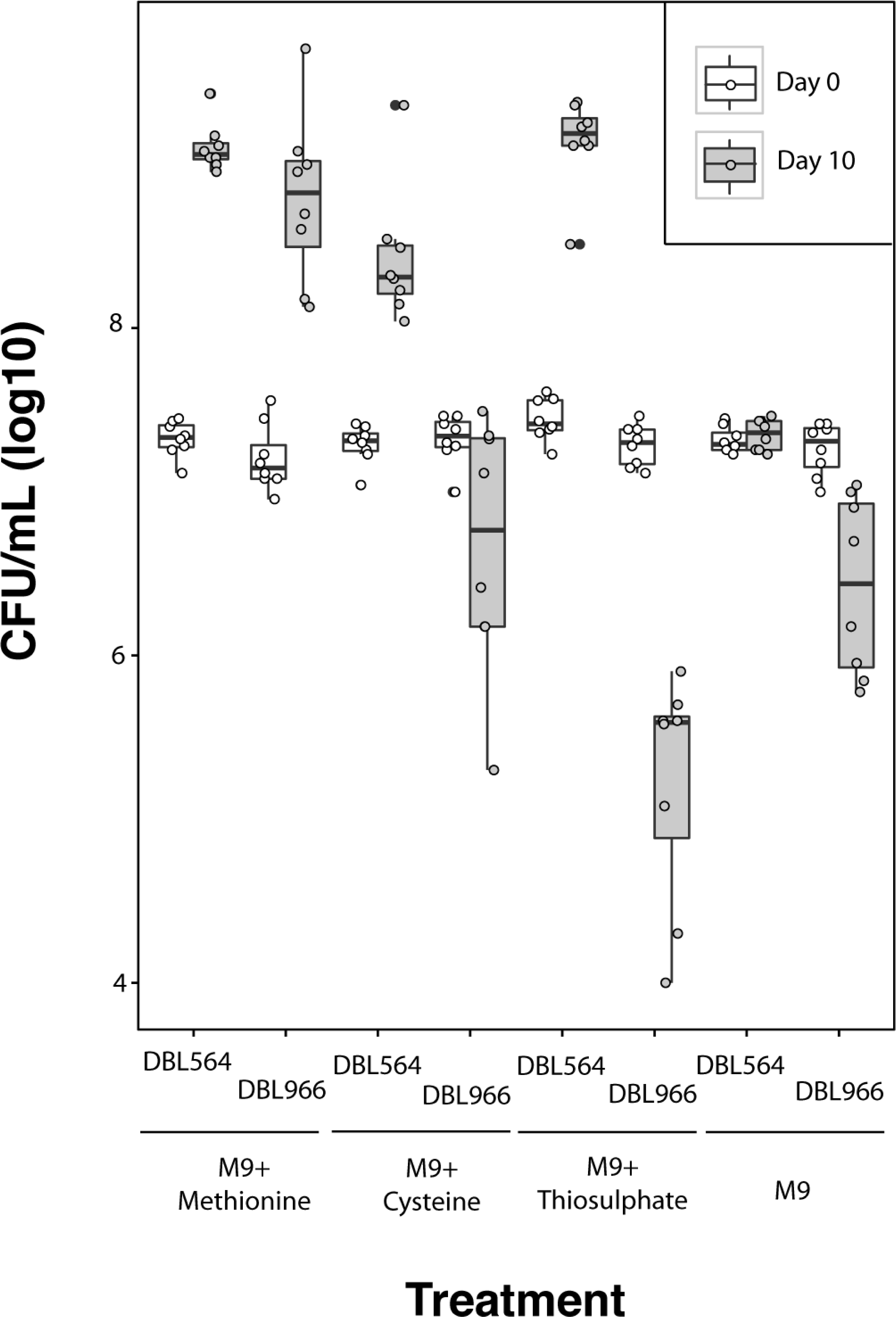
Phenotypic Diversity of Sulfur Utilization Across *Luteibacter* Strains. Plots show each data point for *Luteibacter* sp. 9143 (DBL564) and *Luteibacter* sp. UNCMF366Tsu5.1 (DBL966) in M9 supplemented with a variety of sulfur sources. The same cultures were sampled at day 0 (white plot) and day 10 (grey plot). Methionine and cysteine were supplemented at 100**μ**M. Thiosulfate was supplemented at 10mM. Error bars represent +/- one standard error.

### Sulfur Acquisition by *Luteibacter* Under Natural Growth Conditions

At a minimum, sulfur is required for growth of all known bacteria under all known conditions because it is a main component of an amino acid (methionine) present in all translated proteins (1). Sulfate is one of the most widely available and utilized sulfur sources for terrestrial bacteria, which is highlighted by conservation of the sulfate transport pathway genes *cysAUWP* across many proteobacteria, including xanthomonads (10). That certain clades of *Luteibacter* have altered abilities to metabolize sulfate compared to closely related outgroups like *Xanthomonas* speaks to a distinctive ecological niche for these strains compared to their more widely studied relatives.

For both *Luteibacter* sp. 9143 and UNC366Tsa5.1, sulfur may be acquired through close relationships with fungi and/or plants, rather than from other environmental sources. While sulfur concentrations are thought to be relatively low inside of plants compared to the surrounding environment (1), these strains could be adapted to scavenging free amino acids or other organic sulfur-containing compounds provided by eukaryotic hosts.

Aside from potentially available plant-derived sulfur containing compounds like proteins containing cysteine and methionine, sulpholipids are a main component of plant cells and potentially could be used as a sulfur source by these bacteria (11). *Luteibacter* sp. UNCMF366Tsa5.1 was isolated from the rhizosphere of *Arabidopsis*, but it is possible that this strain was closely associated with fungi in proximity to plant roots (3).

Lastly, we note that *Mycoavidus cysteinexigens*, a symbiont of the fungus *Mortierella elongata*, has lost pathways for the production of cysteine and is therefore a true auxotroph for this amino acid (12). While this is clearly an independent evolutionary event relative to loss of sulfate transport in *Luteibacter* sp., it is intriguing that pathways related to sulfur metabolism have been the targets of repeated modifications in endofungal bacteria. Although evolutionary drivers of this pattern are unclear at present, production of sulfur containing molecules during plant defense responses (13) or by fungi themselves (14) could enable loss of pathways involved in sulfur acquisition for these bacterial lineages.

Alteration of pathways for sulfur metabolism and acquisition due to availability of sources other than sulfate has been demonstrated multiple times in aquatic bacteria (15–17). It is hypothesized that an abundance of dimethylsulphoniopropionate (DMSP), supplied by other marine microbes, enabled the loss of sulfur transporters (18, 19). In some cases, DMSP can be supplied by eukaryotes (e.g., the diatom *Thalassiosira pseudonana,* which forms close relationships with the marine bacteria *Ruegeria pomeroyi*) (20). The assays described here showed that *Luteibacter* sp. 9143 fails to use DMSP as a sulfur source under lab conditions (Figure 2). That sulfate transport has been lost under such a variety of conditions speaks to a potentially significant evolutionary or metabolic cost associated with maintenance of sulfate assimilation pathways.

### Conclusions

The experiments presented here demonstrate that certain *Luteibacter* strains have lost commonly found pathways that enable utilization of sulfate as a sulfur source. Sulfur acquisition by these strains under natural conditions potentially requires access to organic sulfur such as that provided by hosts, which further suggests that symbiosis is a key component of the ecological niche for this subset of strains. These results also suggest that one should be cautious when interpreting differential growth of a focal bacterial strain in rich compared to minimal media as auxotrophy, as strains of interest may have different growth requirements in minimal media than would be commonly assumed based on knowledge of closely related strains.

## Acknowledgements

We thank Kayla Arendt, Kevin Hockett, and Paul Carini for advice with experiments and interpretation. Financial support was provided by the National Science Foundation (NSF IOS-1354219 to D.A.B., A.E.A., and Rachel E. Gallery).

## References

1. Sekowska A, Kung HF, Danchin A. 2000. Sulfur metabolism in Escherichia coli and related bacteria: facts and fiction. J Mol Microbiol Biotechnol 2:145–177.

2. Kertesz MA. 2001. Bacterial transporters for sulfate and organosulfur compounds. Res Microbiol 152:279–290.

3. Lundberg DS, Lebeis SL, Paredes SH, Yourstone S, Gehring J, Malfatti S, Tremblay J, Engelbrektson A, Kunin V, del Rio TG, Edgar RC, Eickhorst T, Ley RE, Hugenholtz P, Tringe SG, Dangl JL. 2012. Defining the core *Arabidopsis thaliana* root microbiome. Nature 488:86–90.

4. Hoffman MT, Arnold AE. 2010. Diverse bacteria inhabit living hyphae of phylogenetically diverse fungal endophytes. Appl Environ Microbiol 76:4063–4075.

5. Hoffman MT, Gunatilaka MK, Wijeratne K, Gunatilaka L, Elizabeth Arnold A. 2013. Endohyphal Bacterium Enhances Production of Indole-3-Acetic Acid by a Foliar Fungal Endophyte. PLoS One 8:e73132.

6. Arendt KR, Hockett KL, Araldi-Brondolo SJ, Baltrus DA, Arnold AE. 2016. Isolation of Endohyphal Bacteria from Foliar Ascomycota and In Vitro Establishment of Their Symbiotic Associations. Appl Environ Microbiol 82:2943–2949.

7. Baltrus DA, Dougherty K, Arendt KR, Huntemann M, Clum A, Pillay M, Palaniappan K, Varghese N, Mikhailova N, Stamatis D, Reddy TBK, Ngan CY, Daum C, Shapiro N, Markowitz V, Ivanova N, Kyrpides N, Woyke T, Arnold AE. 2017. Absence of genome reduction in diverse, facultative endohyphal bacteria. Microb Genom 3:e000101.

8. Markowitz VM, Mavromatis K, Ivanova NN, Chen I-MA, Chu K, Kyrpides NC. 2009. IMG ER: a system for microbial genome annotation expert review and curation. Bioinformatics 25:2271–2278.

9. Bertels F, Silander OK, Pachkov M, Rainey PB, van Nimwegen E. 2014. Automated reconstruction of whole-genome phylogenies from short-sequence reads. Mol Biol Evol 31:1077–1088.

10. Pereira CT, Moutran A, Fessel M, Balan A. 2015. The sulfur/sulfonates transport systems in *Xanthomonas citri pv. citri*. BMC Genomics 16:524.

11. Collier R, Kennedy GY. 1963. The sulpholipids of plants I. The distribution of sulpholipids and phospholipids among the major plant classes. J Mar Biol Assoc U K 43:605.

12. Uehling J, Gryganskyi A, Hameed K, Tschaplinski T, Misztal PK, Wu S, Desirò A, Vande Pol N, Du Z, Zienkiewicz A, Zienkiewicz K, Morin E, Tisserant E, Splivallo R, Hainaut M, Henrissat B, Ohm R, Kuo A, Yan J, Lipzen A, Nolan M, LaButti K, Barry K, Goldstein AH, Labbé J, Schadt C, Tuskan G, Grigoriev I, Martin F, Vilgalys R, Bonito G. 2017. Comparative genomics of Mortierella elongata and its bacterial endosymbiont Mycoavidus cysteinexigens. Environ Microbiol.

13. Bloem E, Haneklaus S, Schnug E. 2014. Milestones in plant sulfur research on sulfur-induced-resistance (SIR) in Europe. Front Plant Sci 5:779.

14. Strobel G, Ford E, Harper JK. March 2007. Pestalotiopsis microsporia isolates and compounds derived therefrom. 7192939. US Patent.

15. Tripp HJ, Kitner JB, Schwalbach MS, Dacey JWH, Wilhelm LJ, Giovannoni SJ. 2008. SAR11 marine bacteria require exogenous reduced sulfur for growth. Nature 452:741–744.

16. Smith DP, Nicora CD, Carini P, Lipton MS, Norbeck AD, Smith RD, Giovannoni SJ. 2016. Proteome Remodeling in Response to Sulfur Limitation in “Candidatus Pelagibacter ubique.” mSystems 1.

17. González JM, Kiene RP, Moran MA. 1999. Transformation of sulfur compounds by an abundant lineage of marine bacteria in the alpha-subclass of the class Proteobacteria. Appl Environ Microbiol 65:3810–3819.

18. Reisch CR, Moran MA, Whitman WB. 2011. Bacterial Catabolism of Dimethylsulfoniopropionate (DMSP). Front Microbiol 2:172.

19. Moran MA, Reisch CR, Kiene RP, Whitman WB. 2012. Genomic insights into bacterial DMSP transformations. Ann Rev Mar Sci 4:523–542.

20. Durham BP, Sharma S, Luo H, Smith CB, Amin SA, Bender SJ, Dearth SP, Van BAS, Campagna SR, Kujawinski EB, Armbrust EV, Moran MA. 2015. Cryptic carbon and sulfur cycling between surface ocean plankton. Proc Natl Acad Sci U S A 112:453–457.

